# A 90K SNP array uncovers inbreeding and cryptic relatedness in an Antarctic fur seal breeding colony

**DOI:** 10.1101/2020.04.01.020123

**Authors:** Emily Humble, Anneke J. Paijmans, Jaume Forcada, Joseph I. Hoffman

## Abstract

High density single nucleotide polymorphism (SNP) arrays allow large numbers of individuals to be rapidly and cost-effectively genotyped at large numbers of genetic markers. However, despite being widely used in studies of humans and domesticated plants and animals, SNP arrays are lacking for most wild organisms. We developed a custom 90K Affymetrix Axiom array for an intensively studied pinniped, the Antarctic fur seal (*Arctocephalus gazella*). SNPs were discovered from a combination of genomic and transcriptomic resources and filtered according to strict criteria. Out of a total of 85,359 SNPs tiled on the array, 75,601 (88.6%) successfully converted and were polymorphic in 274 animals from a breeding colony at Bird Island in South Georgia. Evidence was found for inbreeding, with three genomic inbreeding coefficients being strongly intercorrelated and the proportion of the genome in ROH being non-zero in all individuals. Furthermore, analysis of genomic relatedness coefficients identified multiple second and third order relatives among a sample of ostensibly unrelated individuals. Such “cryptic relatedness” within fur seal breeding colonies may increase the likelihood of consanguinous matings and could therefore have implications for understanding fitness variation and mate choice. Finally, we demonstrate the cross-amplification potential of the array in three related species. Overall, our SNP array will facilitate future studies of Antarctic fur seals and has the potential to serve as a more general resource for the wider pinniped research community.

## INTRODUCTION

Single nucleotide polymorphisms (SNPs) have become one of the most popular genetic markers in evolutionary and conservation biology (Morin *et al*. 2004). They are the most abundant form of genetic variation and in contrast to classical markers such as microsatellites, they can be genotyped on a very large scale (Seeb *et al*. 2011). Consequently, SNPs can provide the resolution needed to address broad reaching questions in ecology, evolution and conservation biology with greater power than was previously possible. In particular, quantitative genetic and gene mapping studies have profited enormously from the power of these markers (Johnston *et al*. 2013; Berenos *et al*. 2014; Barson *et al*. 2015; Gienapp *et al*. 2017).

Two of the most common approaches for genotyping SNPs in non-model organisms are genotyping by sequencing (GBS) methods such as restriction site associated DNA (RAD) sequencing (Hohenlohe *et al*. 2010; Davey *et al*. 2011) and array based methods in which panels of pre-determined polymorphisms are hybridised onto chips by companies such as Affymetrix and Illumina. GBS approaches are capable of genotyping tens of thousands of SNPs and do not necessarily require access to existing genomic resources. However, they generate large amounts of sequence data that require bioinformatic processing, which can be time-consuming and technically challenging (Shafer *et al*. 2017). An additional issue with GBS is that the depth of sequence coverage is not always high enough to call genotypes with high confidence, which leads to high rates of missing data (Chattopadhyay *et al*. 2014; Huang and Knowles 2016; Benjelloun *et al*. 2019). By contrast, array based methods are faster, require minimal technical effort, have low genotyping error rates and high call rates, and can easily be scaled up to very large numbers of individuals. SNP arrays are also flexible, with low density arrays allowing hundreds to thousands of SNPs to be genotyped and high density arrays or “SNP chips” supporting tens of thousands to millions of SNPs (Thaden *et al*.; Shi *et al*. 2012). For these and other reasons, array based genotyping has become the method of choice for many researchers, particularly those working on long-term datasets with access to many individuals.

Until recently, the majority of array based studies of natural populations exploited resources already developed for closely related domestic species such as the BovineSNP50 and OvineSNP50 bead chips (Thaden *et al*.; Shi *et al*. 2012). However, given that cross-species polymorphism declines with increasing phylogenetic distance (Miller *et al*. 2012), custom species-specific arrays are now being developed for several wild species such as great tits (Kim *et al*. 2018), flycatchers (Kawakami *et al*. 2014), house sparrows (Lundregan *et al*. 2018) and polar bears (Malenfant *et al*. 2015). These resources have already provided insights into diverse topics from adaptive divergence and hybridization (Bourret *et al*. 2013; McFarlane *et al*. 2020) through to conservation genomics (Chen *et al*. 2016) and quantitative trait locus mapping (Kim *et al*. 2018). However, high rates of failure are not uncommon with custom arrays, as considerable numbers of SNPs either fail to produce any results at all (i.e. they do not “convert”) or they appear monomorphic and are consequently for most purposes uninformative. Among recent efforts to develop SNP arrays for wild organisms, the proportion of tiled SNPs converting into high quality polymorphic genotyping assays has varied from just over 50% to at most around 80% (van Bers *et al*. 2012; Hagen *et al*. 2013; Kawakami *et al*. 2014; Malenfant *et al*. 2015; Kim *et al*. 2018). Recent studies investigating the causes of assay failure have identified poor SNP genomic context as a major factor, particularly when markers are derived from a transcriptome, and have highlighted the advantages of considering how SNP probe sequences map to a reference assembly (Humble *et al*. 2016a, 2016b). Consequently, incorporating contextual information into SNP filtering pipelines has the potential to substantially improve the success rates of custom arrays.

The Antarctic fur seal (*Arctocephlaus gazella*) is a prime example of a species that would benefit from the development of a SNP array. On Bird Island in South Georgia, a breeding colony of fur seals has been intensively monitored since the 1980s and genetic, phenotypic and life-history data have been collected for around ten thousand animals. This information has provided the foundation for elucidating the species’ mating system (Hoffman *et al*. 2003, 2007), demographic history (Hoffman *et al*. 2011; Paijmans *et al*. 2020) and population status (Forcada and Hoffman 2014). For example, by combining data from nine microsatellites with multi-event mark-recapture models, Forcada and Hoffman (2014) showed that adverse climate effects have led to a 24% decline in the number of breeding females over the past three decades. Alongside this, breeding female heterozygosity has increased by around 8.5% per generation since the early 1990s (Forcada and Hoffman 2014). Together, these patterns are strongly suggestive of increasing viability selection against homozygous individuals, possibly due to inbreeding depression.

To shed light on this phenomenon in fur seals as well as to improve our broader understanding of the mechanisms responsible for inbreeding depression, a shift from using small numbers of microsatellites to many thousands of SNPs is required (Kardos *et al*. 2015). High density datasets of mapped SNPs are capable of estimating inbreeding with extremely high precision because they can capture and measure the genome-wide contribution of runs of homozygosity (ROH), contiguous tracts of homozygous SNPs that occur when individuals inherit two identical by descent (IBD) copies of a chromosomal segment from a common ancestor (Franklin 1977). Indeed, simulation studies have shown that ROH based measures provide more precise estimates of inbreeding than those obtained from pedigrees (Keller *et al*. 2011), which cannot capture variation among individuals due to recombination and Mendelian sampling (Hill and Weir 2011). Furthermore, the length distribution of ROH can shed light on whether inbreeding is the result of matings between relatives in recent generations or in the distant past (Thompson 2013). This is because the length of an IBD segment is determined by the number of generations between the inbred individual and the most recent common ancestor carrying the two homologous copies of that IBD segment. For these reasons, quantifying ROH is becoming the method of choice among researchers interested in inbreeding and inbreeding depression (Kardos *et al*. 2017; Grossen *et al*. 2018; van der Valk *et al*. 2020).

As well as improving estimates of inbreeding, genome-wide marker panels have also made it possible to calculate precise measures of relatedness, something that has traditionally been restricted to populations for which a pedigree is available (Santure *et al*. 2010; Huisman 2017). Understanding how animals are related is of fundamental importance to many aspects of evolutionary and conservation biology, from understanding patterns and mechanisms of mate choice (Foerster *et al*. 2006; Blyton *et al*. 2016; Tuni *et al*. 2019) to making informed pairing decisions in conservation breeding programmes (Galla *et al*. in press). As high quality, multi-generational pedigrees are not available for most wild populations, the possibility of using genomic data for deriving relatedness estimates therefore provides many additional research opportunities.

This paper describes the development of a 90K Affymetrix Axiom genotyping array for the Antarctic fur seal. As our longer-term aims are to investigate the mechanism(s) behind the population decline as well as more generally to explore the genetic architecture of fitness-related traits, we developed a genome-wide panel of nuclear SNPs based on RAD sequencing data from a recent study (Humble et al 2018). We additionally made use of another desirable property of SNP arrays, the possibility of incorporating candidate gene markers, by tiling over ten thousand polymorphisms from a transcriptome assembly (Humble *et al*. 2016b) together with a handful of SNPs from the major histocompatibility complex (MHC), a group of genes constituting arguably the most important component of the vertebrate immune system (Sommer 2005). Finally, we attempted to maximise the overall genotyping success of the array by subjecting all discovered SNPs to a strict prioritisation scheme that incorporated multiple sources of information including the genomic context of each locus. We genotyped 288 samples, primarily from Antarctic fur seals but also including three additional pinniped species, to assess the performance of the SNP array, to quantify inbreeding and to explore patterns of relatedness among individuals.

## MATERIALS AND METHODS

### Genomic SNP discovery

Genome-wide distributed nuclear SNPs were discovered using RAD sequencing as described by Humble *et al*. (2018). Briefly, tissue samples from 83 individuals were collected from the main breeding colonies across the species range: Bird Island, South Georgia (*n* = 57), Cape Shirreff in the South Shetlands (*n* = 6), Bouvetøya (*n* = 5), Îles Kerguelen (*n* = 5), Heard Island (*n* = 5) and Macquarie Island (*n* = 5). RAD libraries were prepared using a protocol with minor modifications as described in Matsuzaki *et al*. (2004). Read quality was assessed using FastQC v0.112 and the sequences were trimmed to 225 bp and demultiplexed using *process_radtags* in STACKS v1.41 (Catchen *et al*. 2013). To identify SNPs to include on the array, we followed GATK’s best practices workflow (Poplin *et al*. 2017) using the Antarctic fur seal genome v1.2 as a reference (Humble *et al*. 2016a). The resulting SNP dataset was filtered to include only biallelic SNPs using bcftools (Li 2011).

We then applied a set of initial quality filters using vcftools (Danecek *et al*. 2011) to filter out low quality SNPs from our dataset. Specifically, we removed genotypes with a depth of coverage of less than five or greater than 18 to minimise spurious SNP calls due to low coverage or repetitive genomic regions. We also removed SNPs with minor allele frequencies (MAF) below 0.05 and with a genotyping rate below 60%. Next, to prepare the remaining loci for array design, we filtered out SNPs with insufficient flanking sequences by identifying and removing those less than 35 bp away from the start or end of a scaffold. We then collated a list of probe sequences for the remaining SNPs by extracting their 35 bp flanking sequences from the Antarctic fur seal reference genome using the BEDTOOLS command getfasta (Quinlan and Hall 2010).

### Transcriptomic SNP discovery and annotation

In order to allow polymorphisms residing within expressed genes to be genotyped on the array, we included SNPs discovered from the Antarctic fur seal transcriptome in our list of probe sequences. The transcriptome sequencing, assembly and SNP detection process is fully described in (Humble *et al*. 2016b). In brief, testis, heart, spleen, intestine, kidney and lung samples were obtained from nine Antarctic fur seals that died of natural causes at Bird Island, South Georgia. Skin samples were additionally collected from 12 individuals from the same locality. The transcriptome was assembled in multiple iterations using 454 and Illumina sequence data from three different cDNA libraries (Hoffman 2011; Hoffman *et al*. 2013b; Humble *et al*. 2018). SNPs were then discovered using four separate genotype callers and reduced to a consensus subset that was identified by all methods. Loci with sufficient flanking sequences for probe design, and which had been assigned appropriate quality scores by Affymetrix in our previous study, were retained for array design.

Putative functions were assigned to the transcriptomic SNPs by BLASTing the transcripts against the SwissProt, Trembl and non-redundant blast databases using BLASTx v2.2.30 with an e-value cutoff of 1e_-4_. We then used the total_annotation.py script provided by the Fool’s Guide to RNAseq (De Wit *et al*. 2012) to combine all BLAST results, download Uniprot flat files and extract Gene Ontology (GO) categories. To track the number of SNPs with putative immune, growth and metabolism functions throughout the array design process, we flagged all SNPs residing within transcripts associated with the annotation terms described in Table S1.

### Pre-validated and MHC-derived SNPs

We also added to our list of probe sequences a further set of SNPs that were previously demonstrated to be polymorphic in the study colony. These included 40 SNPs derived from RAD sequencing data that were validated using Sanger sequencing (Humble *et al*. 2018), 102 transcriptomic SNPs that were validated using Illumina’s GoldenGate assay (Hoffman *et al*. 2012) and 173 cross-amplified SNPs from the Canine HD Bead chip that were previously shown to be polymorphic in 24 Antarctic fur seals (Hoffman *et al*. 2013a). In addition to these, we included a further six SNPs that were recently discovered from the second exon of the Antarctic fur seal MHC DQBII locus based on Illumina MiSeq data from 82 Antarctic fur seals (Ottensmann and Hoffman, unpublished data).

### SNP selection

We took our combined list of probe sequences, comprising genomic and transcriptomic SNPs together with pre-validated and MHC-derived SNPs, and evaluated their suitability for inclusion on an Affymetrix Axiom SNP genotyping array. First, we assessed the genomic context of each SNP by blasting their flanking sequences against the fur seal reference genome using BLASTN v2.2.30 with an e-value threshold of 1e_-12_. We then determined the total number of mappings and the alignment length of the top BLAST hit. Finally, all of the probe sequences were sent to Affymetrix who assigned recommendations to each SNP using an *in silico* evaluation tool. This tool considers probe sequence characteristics such as GC content and flanking sequence duplication and calculates a probability of successfully converting into a genotyping assay for each locus. We then prioritised a list of SNPs to be included on the array based on the following criteria:

i. Priority one was assigned to SNPs with an Affymetrix recommendation of “recommended” in either the forward or reverse direction, that mapped uniquely and completely to the reference genome and that were neither an A/T nor a C/G SNP, as these require twice the number of probes. We also assigned priority one status to all pre-validated and MHC-derived SNPs regardless of their Affymetrix design scores.
ii. Priority two status was assigned to the remaining loci if they had a “neutral” recommendation by Affymetrix in either the forward or reverse direction, mapped uniquely and completely to the reference genome, were neither an A/T nor a C/G SNP and had no secondary SNPs present within the flanking sequence.
iii. Priority three status was assigned to any remaining RAD loci with an Affymetrix recommendation of “recommended” in either the forward or reverse direction, that mapped to no more than two different locations in the reference genome, that were neither an A/T nor a C/G SNP and had a MAF of at least 0.017 in South Georgia (equivalent to the minor allele having been found in at least two individuals in the discovery pool for this population). The latter filter was to prioritise SNPs that were polymorphic in our study population.
iv. Priority four status was assigned to the remaining RAD loci with an Affymetrix recommendation of “recommended” in either the forward or reverse direction, that mapped to no more than three different locations in the reference genome and that were neither an A/T nor a C/G SNP.
v. Priority five status was assigned to any high-quality A/T or C/G SNPs that were assigned an Affymetrix recommendation of “recommended” in either the forward or reverse direction and that mapped uniquely and completely to the reference genome.
vi. Priority six status was assigned to all remaining RAD loci with a “neutral” recommendation in either the forward or reverse strand, that mapped to no more than two different locations in the reference genome, that were neither an A/T nor a C/G SNP and that had no secondary SNPs present within the flanking sequence.
vii. Priority seven status was assigned to all remaining RAD SNPs with neutral recommendations for either the forward or reverse strand.

Any SNPs remaining after these prioritisation steps were assigned a priority of zero and were no longer considered for array design. After determining the priority of each SNP, we then thinned the dataset so that all RAD derived SNPs with a priority greater than or equal to three were at least 1 kb from the next adjacent SNP, and all SNPs with a priority of one or two were at least 100 bp apart. Finally, we removed 289 duplicate SNPs that were discovered by more than one approach. The final set of 87,608 SNPs was submitted to Afymettrix for Axiom myDesign chip manufacture.

### Genotyping

To assess the performance of the genotyping array, a total of 288 samples on three 96 well plates were genotyped on a Gene Titan platform by the Beijing Genomics Institute (BGI). To estimate the overall genotyping error rate, a single fur seal individual was genotyped three times, once on each plate. The majority of samples (*n* = 276) were collected from Antarctic fur seals at Bird Island, South Georgia as part of a long-term monitoring study conducted by the British Antarctic Survey. These were made up of females born between 1984 and 2016 and included 53 mother-offspring pairs. Additionally, we evaluated cross-species amplification by genotyping four samples each of three pinniped species including one phocid (the Grey seal, *Halichoerus grypus*) and two otariids (the Steller’s sea lion, *Eumetopias jubatus*, and the Galápagos sea lion, *Zalophus wollebaeki*). DNA was extracted using a standard phenol-chloroform protocol (Sambrook and Russell 2006) and quantified using PicoGreen® on a TECAN Infinite® 200 PRO plate reader. A total of 271 samples had DNA concentrations above the manufacturer’s recommendation of 50 ng/µl. The remaining 15 samples had DNA concentrations between 40 and 50 ng/µl (*n* = 7) or between 20 and 40 ng/µl (*n* = 8). These were included to evaluate how samples with suboptimal DNA concentrations would perform on the array.

The resulting genotype data were analysed using Affymetrix Power Tools (APT) command line software. We applied two workflows to the data, the first to assess the performance of the array in the Antarctic fur seal, and the second to quantify rates of cross-species amplification. For the former, we excluded samples belonging to the other three pinniped species so that their inclusion did not impact overall cluster quality, and then filtered out samples with dish QC scores less than 0.82 and with call rates below 97%. For the latter, we excluded samples with dish QC scores below 0.82 but did not filter on the basis of call rate in order to retain as many samples from the other pinniped species as possible.

Genotyping was conducted for both datasets using the apt_genotype_axiom function in APT, with quality metrics and classifications being assigned to individual SNPs using the Ps_Metrics and Ps_Classification functions respectively. We then used the OTV_Caller function in the SNPolisher R package to recover SNPs that were originally classified as “off-target variants”. The resulting output was then re-classified using the APT functions Ps_Metrics and Ps_Classification. To estimate the genotyping error rate, we quantified the probability at each typed locus of both alleles being IBD between replicate samples using the Z2 score output of the --genome command in PLINK v1.9 (Purcell *et al*. 2007).

### Inbreeding

The genomic data were subsequently used to estimate levels of inbreeding in our study population. In order to generate a high-quality dataset with minimal missing data, we selected all of the polymorphic SNPs and then used PLINK to retain loci with a genotyping rate of over 90%, MAF > 0.01 and that conformed to Hardy-Weinberg equilibrium with a *p-*value threshold of 0.001. Using the resulting dataset of 74,261 SNPs genotyped in 272 individuals, we calculated three genomic estimates of inbreeding for each individual: standardised multi-locus heterozygosity (sMLH), a measure based on the correlation of uniting gametes (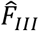), and the proportion of the genome in ROH (*F*_ROH_). sMLH was calculated using the R package inbreedR (Stoffel *et al*. 2016) and 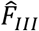 was calculated using the --ibc function in GCTA (Yang *et al*. 2011).

To calculate *F*_ROH_, we first identified regions of the genome in ROH using the --homozyg function in PLINK with a sliding window of 20 SNPs (--homozyg-window-snp 20). A window was defined as homozygous when it contained no more than one heterozygous site (--homozyg-window-het 1) and no more than five missing sites (--homozyg-window-missing 5). If at least 5% of all windows containing a given SNP were defined as homozygous, the SNP was presumed to lie within a homozygous segment (--homozyg-window-threshold 0.05). Homozygous segments were then called as ROH when they contained at least 20 SNPs (--homozyg-snp 20) and no more than one heterozygous site (--homozyg-het 1). Furthermore, to ensure that incomplete marker information did not bias ROH detection, segments were only called as ROH when they contained at least one SNP per 100 kb (--homozyg-density 100) and were at least one Mb in length (-- homozyg-kb 1000). If two SNPs within an ROH segment were further than 1000 kb apart, the ROH was split into two segments (--homozyg-gap 1000). The proportion of the genome in ROH (*F*_ROH_) was then calculated as the sum of the detected ROH lengths for each individual over the total assembly length (2.3 Gb). In addition to these inbreeding estimators, we also quantified the extent of identity disequilibrium using the measure *g*_2_ in inbreedR.

### Relatedness

Next, we used the SNP dataset to infer patterns of relatedness among the Antarctic fur seal individuals. For this analysis, we pruned the dataset of polymorphic SNPs for linkage disequilibrium using the --indep function in PLINK. We used a sliding window of 50 SNPs, a step size of 5 SNPs and removed all variants in a window above a variance inflation factor threshold of 2, corresponding to *r*^*2*^ = 0.5. We then excluded SNPs that deviated significantly from HWE as described above. Finally, in order to retain a subset of SNPs that contained as much information as possible for inferring relationships among individuals, we filtered out loci with MAF below 0.3 and that had been called in fewer than 90% of individuals. Based on the resulting dataset of 6,579 SNPs, we quantified relatedness among all 272 individuals using the --genome function in PLINK. Specifically, we used the measure PI_HAT, which estimates the overall proportion of the genome that is identical by descent (IBD) between pairs of individuals, as well as Z scores, which reflect the probability of sharing zero, one or two alleles IBD. The latter probabilities depend directly on relatedness and therefore provide a more precise method for inferring the type of relationship between individuals (Galván-Femenía *et al*. 2017).

In addition to measures of IBD sharing, we used the R package sequoia version 1.3.5 (Huisman 2017) to assign kinship categories. We first ran an initial iteration of parentage assignment to identify duplicate individuals as well as loci with Mendelian errors by setting *MaxSibIter* to zero. This correctly identified the sample genotyped in triplicate as well as 44 SNPs with Mendelian errors, which were removed from the dataset. We then ran a second iteration of sequoia to assign siblings and second degree relationships by setting *MaxSibIter* to five. For both iterations, birth year information was provided using the *LifeHist* parameter.

### Cross-species amplification potential

Finally, we investigated the cross-amplification potential of the array by quantifying the number of markers that could be successfully called in the grey seal, the Galápagos sea lion and the Steller’s sea lion using the --missing function in PLINK. We furthermore quantified the proportion of called SNPs that were polymorphic in each species.

### Data availability

Flanking sequences and accompanying metadata for all of the SNPs that were printed on the array have been deposited on the European Variation Archive (ENA, https://www.ebi.ac.uk/eva) under study accession number XXX. Code for the analyses are available at https://github.com/elhumble/Agaz_90K_workflow_2018. Supplementary material and SNP genotypes are available via Figshare: XXX.

## RESULTS

### Overview

We discovered SNPs from a combination of genomic and transcriptomic resources, applied appropriate downstream filters, and then selected the most suitable loci for tiling on a custom Antarctic fur seal SNP array according to the priority scheme described in the Materials and methods. Figure 1 summarises the design and implementation of the array including the number of SNPs retained at each step of the selection procedure and the genotyping outcomes for different types and priority categories of SNP.

**Figure 1:**
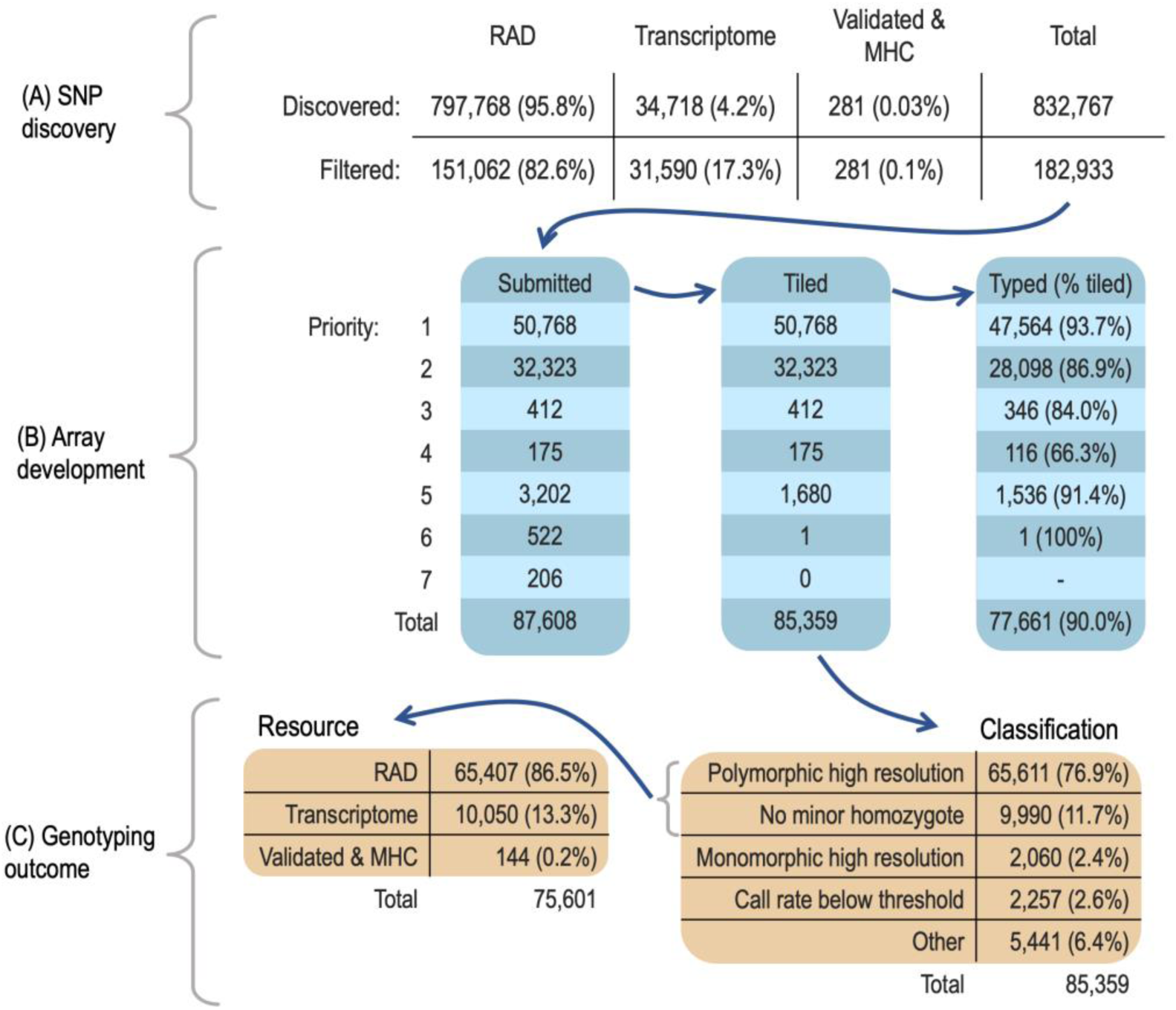
Flow diagram outlining the number of SNPs at each step of the array development pipeline. (A) Numbers of SNPs discovered, filtered and submitted for array design. (B) Numbers of submitted, tiled and genotyped SNPs in priority categories one to seven. (C) Classification outcomes of genotyped SNPs and the breakdown of resource categories for polymorphic SNPs.

### SNP discovery, filtering and array design

Briefly, RAD sequencing data from 83 individuals were used to call a total of 797,768 biallelic SNPs with GATK’s best practices workflow (Humble *et al*. 2018). Downstream filtering for depth of coverage, MAF and genotyping rate resulted in a total of 151,063 SNPs, of which 151,062 had sufficient flanking sequences for probe design. A further 34,718 high quality SNPs were discovered from the Antarctic fur seal transcriptome, of which 32,727 had sufficient flanking sequences for probe design and 31,590 had appropriate Affymetrix quality scores (Humble *et al*. 2016b). Combining the RAD and transcriptomic SNPs resulted in a total of 182,652 loci. These were pooled together with 275 pre-validated SNPs and six SNPs from the MHC to produce a total of 182,933 markers to be considered for array development (Figure 1A). To select the most suitable SNPs for array design, we considered the type of SNP, genomic context, Affymetrix design score metrics, pre-validation status, MAF and spacing of each locus. Based on this information, a total of 87,608 SNPs were assigned to priority categories one to seven and were therefore sent to Affymetrix for printing. Of these, 85,359 (97%) were successfully tiled on the array, of which 59.5% belonged to the highest priority category (Figure 1B).

### Performance of the array

To evaluate the performance of the array, we genotyped a total of 276 Antarctic fur seal individuals across three microtiter plates. To provide a positive control and for genotyping error rate estimation, one of these individuals was genotyped in triplicate, once on each plate. Consequently, the total number of Antarctic fur seal samples genotyped on the array was 278. Four of these samples either failed quality control (*n* = 1) or fell below the call rate threshold of 97% (*n* = 3) and were therefore removed from the dataset. The remaining 274 samples were successfully genotyped at 77,661 SNPs, corresponding to an overall success rate of 90.0% (Figure 1B). These included 163 SNPs that were recovered after having been originally classified as “off-target variants”. The error rate determined from the individual genotyped in triplicate was low at 0.004 per locus.

To evaluate the success of our selection criteria, conversion rates (defined as the proportion of SNPs yielding high quality genotypes) were quantified separately for each priority category. SNPs assigned to priority categories one, five and six had conversion rates in excess of 90% (Figure 1B). Conversion rates were slightly lower (≥ 80%) for priority two and three SNPs, while loci assigned to priority category four had the lowest overall conversion rate of 66.3%. Contrary to expectations, we did not find that pre-validated SNPs had higher conversion rates than SNPs that were not validated in advance. Instead, SNPs from the canine HD Bead chip were actually less likely to convert than non-validated SNPs (58 / 173 [33.5%]; Fisher’s exact test: odds ratio = 0.05, 95% CI = 0.04–0.07, *p* < 0.05). After excluding these loci, the overall conversion rate of pre-validated SNPs did not differ significantly from that of the remaining ones (130 / 142 [91.5%] versus 77,473 / 85,044 [91.1%]; Fisher’s exact test: odds ratio = 1.08, 95% CI = 0.60–2.15, *p* = 0.88).

Overall, no relationship was found between genotyping success, expressed as the call rate per sample, and DNA concentration (slope = −3.36, t = −1.09, df = 278, *P* = 0.27, Figure S1). All fifteen of the samples submitted for genotyping with DNA concentrations below the manufacturer’s recommendation of 50 ng/µl had call rates above 98%, whereas the four samples that were excluded from the final dataset on the basis of suboptimal quality or call rates had DNA concentrations above 50 ng/µl.

### Levels of polymorphism

A total of 75,601 SNPs were polymorphic, equivalent to 88.6% of the tiled loci or 97.3% of the successfully converted loci (Figure 1C). The final dataset of polymorphic loci comprised 65,407 SNPs discovered from the RAD sequencing data, 49 SNPs that were cross-amplified from the canine HD bead array and 10,142 transcriptomic SNPs, which include 92 pre-validated SNPs and three SNPs from the MHC. The loci originating from the RAD data were distributed across 835 genomic scaffolds and had a mean spacing of 35.5 kb (range = 0.02–3306.6 kb, Figure S2). The transcriptomic loci included 1,137 SNPs residing within genes with annotations relating to immunity plus 1,310 SNPs residing in genes with annotations involving metabolism and growth.

Focusing on the polymorphic loci, we investigated patterns of genetic variability by deriving minor allele frequency (MAF) distributions separately for the RAD and transcriptomic SNPs. We also examined the correspondence between variability inferred from animals genotyped on the array (“empirical MAF”) and variability inferred from the original genomic and transcriptomic resources (“*in silico* MAF”). Empirical MAF was left skewed among the RAD SNPs (Figure 2A, mean = 0.19 +/- 0.13 SD) whereas the transcriptomic SNPs were more evenly distributed across the site frequency spectrum (mean = 0.22 +/- 0.14 SD). The empirical MAF distributions of both classes of marker also extended down to zero (Figure 2A), whereas the corresponding *in silico* values were truncated to 0.05 due to filters applied during the SNP discovery process. A strong positive association was found between empirical and *in silico* MAF for the RAD SNPs (Figure 2B, correlation coefficient = 0.90) but this was somewhat weaker for the transcriptomic SNPs (Figure 2C, correlation coefficient = 0.43).

**Figure 2:**
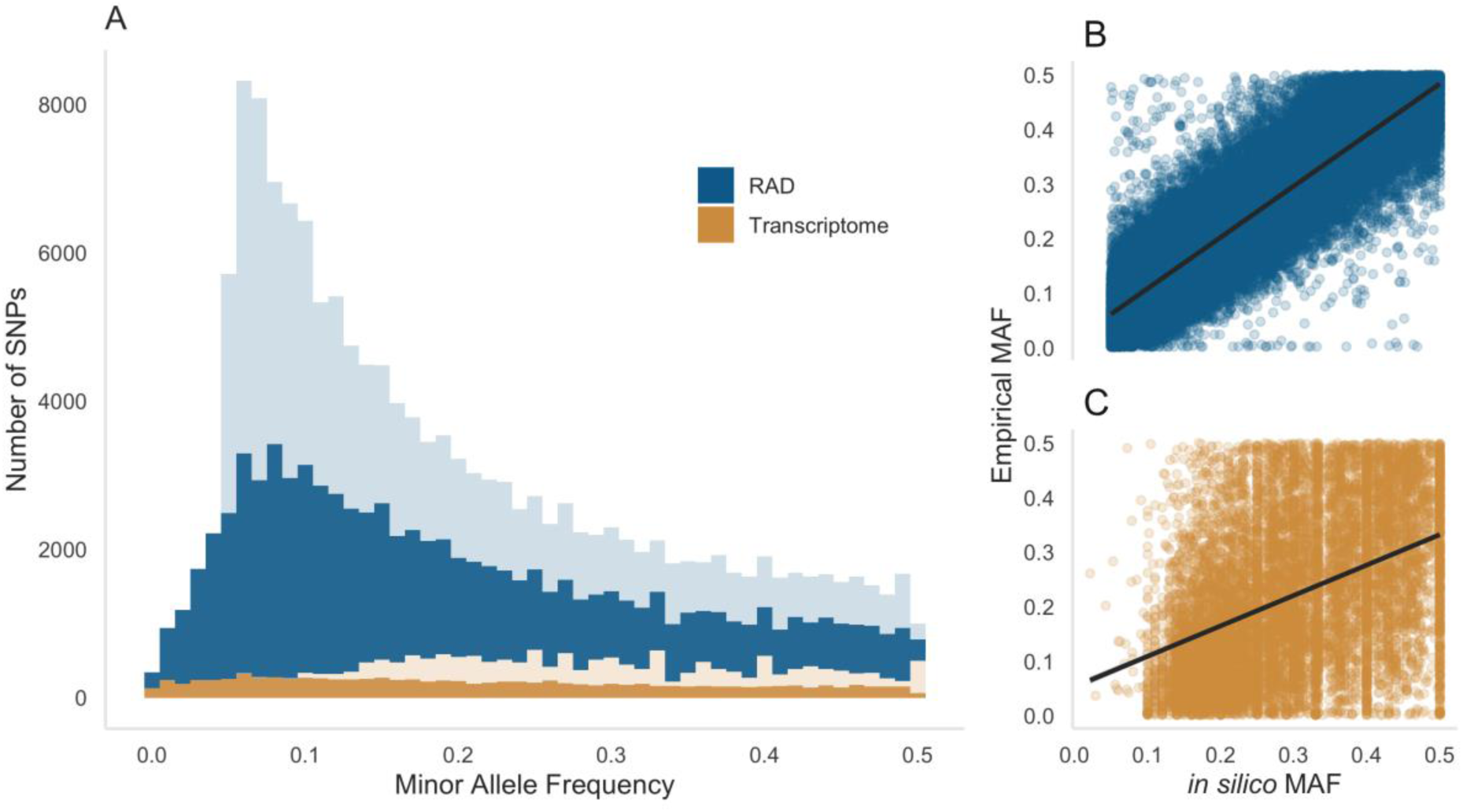
Inferred levels of SNP variability in Antarctic fur seals. (A) Minor allele frequency (MAF) distributions of RAD and transcriptomic SNPs. Dark colours represent empirical MAF and light colours represent *in silico* MAF. Panels on the right-hand side show the strength of association between empirical and *in silico* MAF for (B) the RAD and (C) the transcriptomic SNPs.

### Inbreeding

Inbreeding was investigated using two complementary approaches. First, we quantified identity disequilibrium using the measure *g*^2^, which differed significantly from zero (0.00012, bootstrap 95% confidence interval = 0.000099–0.000149, *p* = 0.001). Second, we calculated for each individual (i) sMLH, an estimate of genome-wide heterozygosity; (ii) 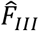, a genomic inbreeding estimator based on the correlation of uniting gametes; and (iii) *F*_ROH_, an estimate of the proportion of the genome in ROH. All three genomic inbreeding measures were intercorrelated (*r* = 0.62– 0.87, Figure 3B–D) and *F*_ROH_ was non-zero for every individual (Figure 3A, mean = 0.06, range = 0.03–0.08). The length distribution of ROH ranged from one to 22 Mb, with short ROH (< 5 Mb) making up a larger proportion of the genome than medium or long ROH (≥ 5 Mb) (Figure 4A). In particular, ROH < 5Mb had a total median length of 106 Mb whilst long ROH ≥ 5Mb had a total median length of 19.1 Mb. ROH longer than 20 Mb were only observed in four individuals (Figure 4B).

**Figure 3:**
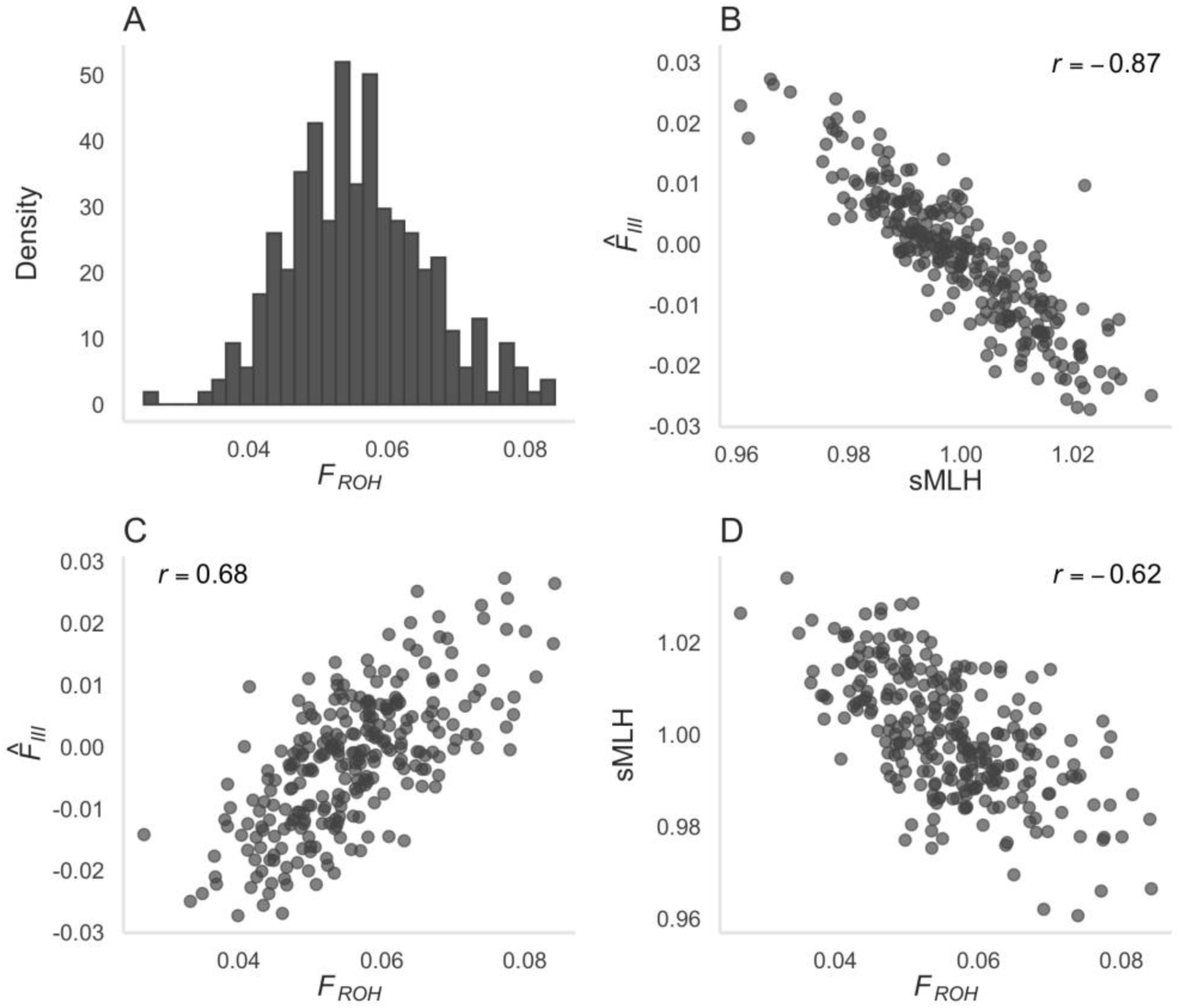
(A) Distribution of *F*_ROH_ values (the estimated proportion of the genome in ROH) for 272 Antarctic fur seals genotyped at 74,261 SNPs; (B–D) Pairwise correlations between the genomic inbreeding coefficients sMLH, 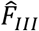 and *F*_ROH_. See the Materials and methods for further details.

**Figure 4:**
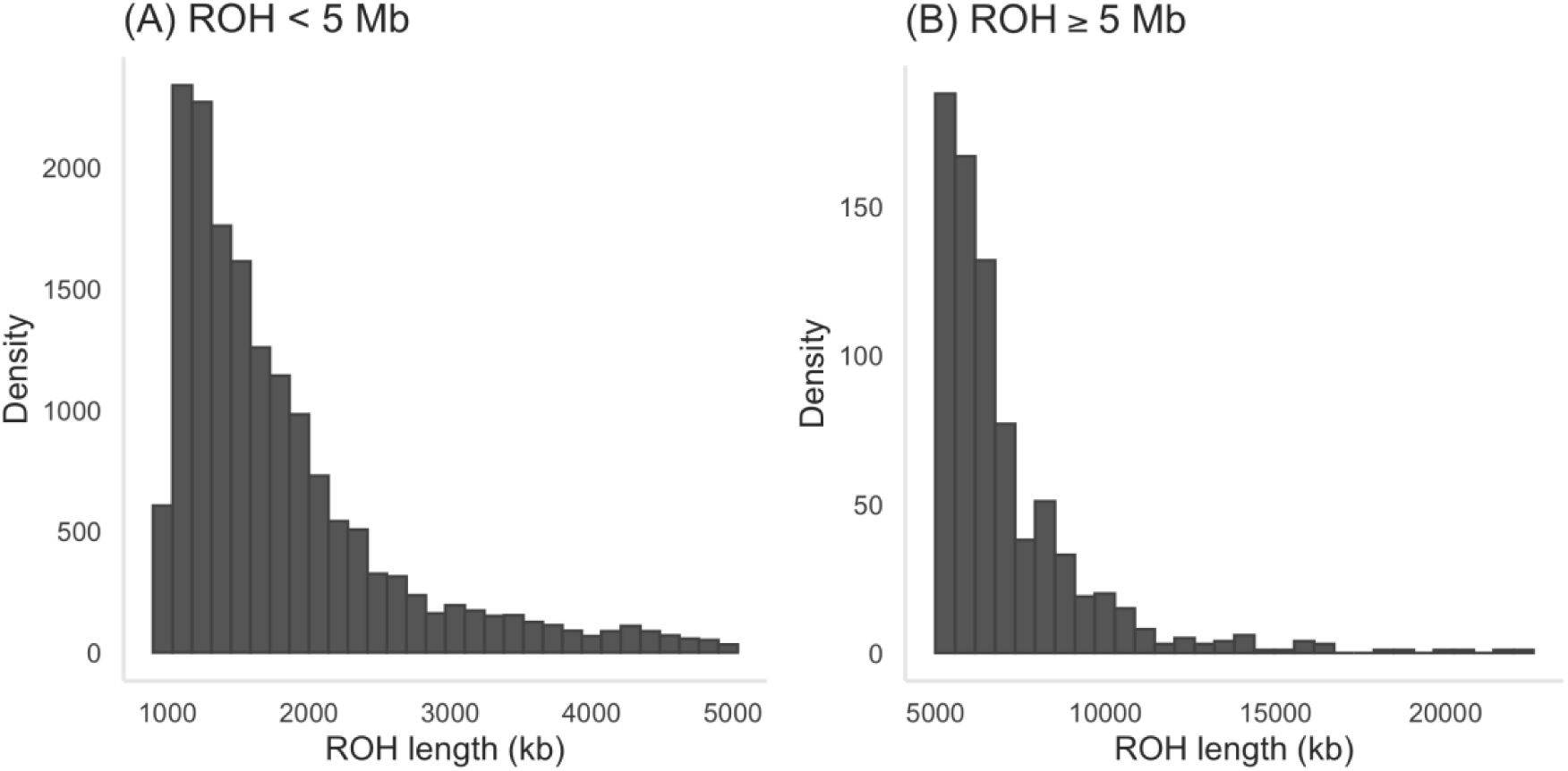
Length distributions of ROH in 272 Antarctic fur seals genotyped at 74,261 SNPs. (A) ROH segments shorter than 5 Mb and therefore due to more recent inbreeding; and (B) ROH segments longer than or equal to 5 Mb and therefore due to inbreeding in the more distant past.

### Relatedness structure

In order to infer patterns of relatedness within our dataset, we analysed a maximally informative dataset of 6,579 polymorphic SNPs genotyped in 272 individuals. A narrow peak of relatedness was present at one, which corresponds to a single individual that was genotyped in triplicate (Figure 5A). A peak was also present at around 0.5, corresponding to 52 mother-offspring pairs in the dataset (Figure 5A). These comprised 48 pairs identified on the basis of field records plus four mother-offspring pairs that were not previously known to be filial pairs. We also identified five pairs of animals that were incorrectly assigned as mother-offspring pairs in the field. These had relatedness values of between zero and 0.23 as opposed to the expectation of around 0.5. The majority of other animals were unrelated, with 99% of pairwise comparisons yielding genomic relatedness coefficients of less than 0.1. However, 64 pairs of individuals had genomic relatedness coefficients of between 0.1 and 0.3, consistent with the presence of multiple second and third order relatives in the study population. We have termed these individuals “cryptic relatives” as they were not previously known to be related.

**Figure 5:**
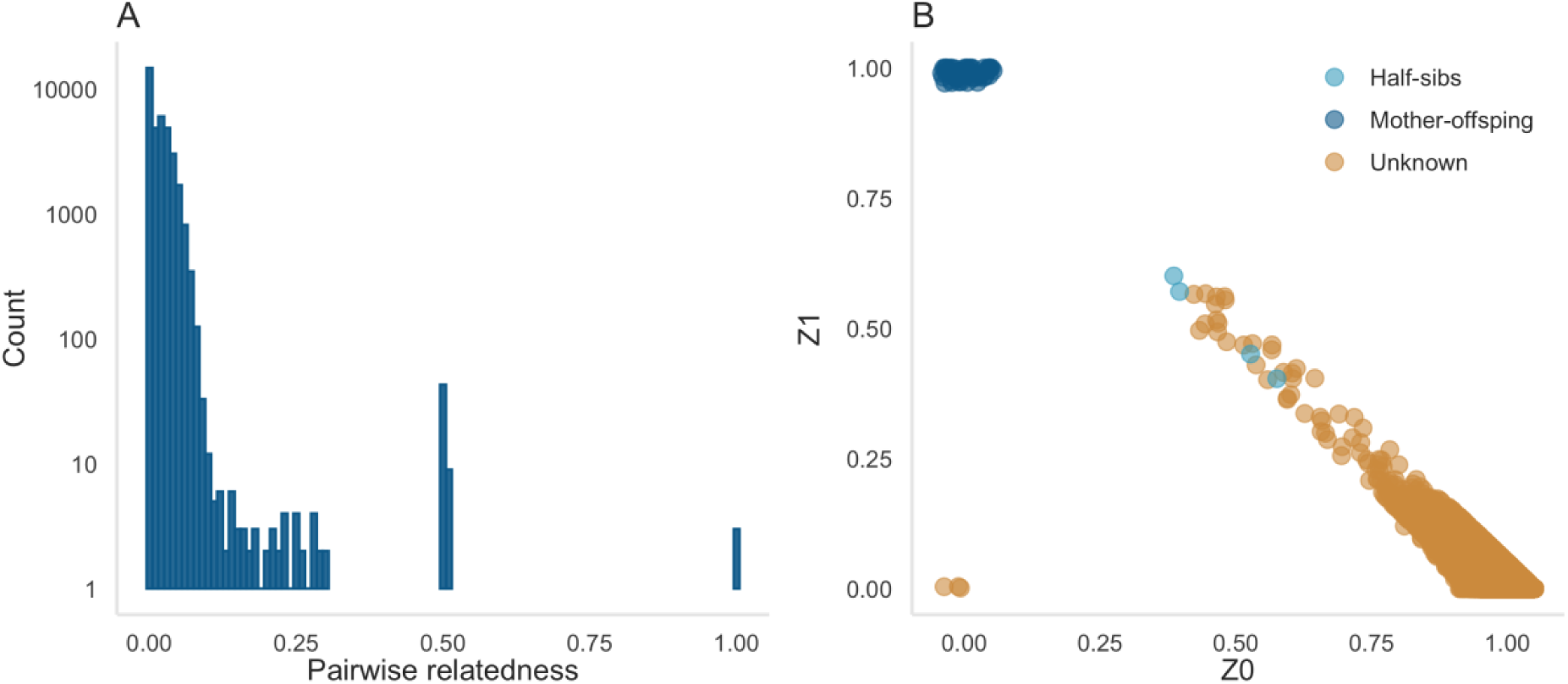
(A) Distribution of genomic relatedness values among all possible pairwise comparisons of Antarctic fur seal individuals in our dataset. Relatedness was quantified as the proportion of the genome identical by descent (IBD) between each pair of individuals based on a dataset of 6,579 maximally informative SNPs (see Materials and methods for details); (B) The probability of sharing zero IBD alleles (Z0) versus the probability of sharing one IBD allele (Z1) for all individual pairwise comparisons. The expectations for specific classes of relative are as follows: Unrelated: Z0 = 1, Z1 = 0, Z2 = 0; Parent offspring: Z0 = 0, Z1 = 1, Z2 = 0; Full siblings: Z0 = 0.25, Z1 = 0.5, Z2 = 0.25; Half siblings, avuncular relationships and grandparents-grandchildren: Z0 = 0.5, Z1 = 0.5, Z2 = 0; First cousins: Z0 = 0.75, Z1 = 0.25, Z2 = 0; Duplicate samples: Z0 = 0, Z1 = 0, Z2 = 1. A small amount of variation (0.05) was applied to the location of each data point to avoid over-plotting and improve interpretation.

To investigate further, we calculated Z scores, which reflect the probability of pairs of individuals sharing zero (Z0), one (Z1) or two (Z2) IBD alleles (Figure 5B). Z scores facilitate more in-depth interpretation of our data as they can be compared against expectations for different categories of relative, which are given in the legend of Figure 5. We plotted Z0 against Z1 for all possible pairwise combinations of individuals in our sample (Figure 5B). Once again, known relationships could be identified, with the triplicate genotypes of the positive control (Z0 = 0 / Z1 = 0) clustering together in the bottom left of the figure and mother-offspring pairs (Z0 = 0 / Z1 = 1) clustering together in the upper left corner. The majority of individuals were unrelated (Z0 = ∼1 / Z1 = ∼0) and therefore clustered in the bottom right corner of Figure 5B. However, a long tail of progressively more related individuals extended along the hypotenuse, with third order relatives clustering around Z0 = 0.75 / Z1 = 0.25, and second order relatives clustering around Z0 = 0.5 / Z1 = 0.5. Full siblings (Z0 = 0.25 / Z1 = 0.5) were notably absent from the dataset.

To delve into more detail, we used the pedigree reconstruction package sequoia to assign kinship categories based on a combination of known relationships and genomic data. Sequoia identified the same mother-offspring pairs as described above. Of the 64 pairs of individuals with relatedness coefficients between 0.1 and 0.3, sequoia assigned paternal half-sib status to four (depicted as light blue points in Figure 5B). Three of these pairs comprised two pups born to different females in successive years, whereas the fourth pair comprised a pup and a breeding female of unknown age that were sampled seven years apart.

### Cross-species amplification

Finally, we investigated the cross-amplification potential of the array by genotyping twelve additional samples belonging to three different pinniped species. All four grey seal samples failed to pass the quality control step and were not considered further. For the Galápagos and Steller’s sea lions, the mean number of SNPs successfully called across individuals was 73,922 (range = 73,109– 74,611) and 74,130 (range= 73,164– 74,583) respectively. This is equivalent to a call rate of 96.2% for the Galápagos sea lion and 96.5% for the Steller’s sea lion. Of those SNPs that could be genotyped, 4,480 (6.1%) were polymorphic in the Galápagos sea lion and 4,191 (5.7%) were polymorphic in the Steller’s sea lion.

## DISCUSSION

We developed a custom 90K SNP array for the Antarctic fur seal. Our efforts to prioritise high quality SNPs for tiling on the array resulted in a relatively high conversion rate, with 88.5% of the tiled loci generating readily interpretable and polymorphic genotypes. Furthermore, call rates were in excess of 99% for the majority of individuals and the genotyping error rate was low at 0.004 per reaction. Analysis of data from 276 fur seals genotyped at 75,601 polymorphic SNPs provided new insights into inbreeding, through measures of ROH, and provided a more refined picture of the relatedness structure of the population. Although our dataset of individuals is still modest, this study provides a first impression of the promise of this array for population genomic studies of an emerging model marine mammal species.

### Design and performance of the array

Designing SNP arrays for non-model species is non-trivial and conversion rates are not always as high as expected (Helyar *et al*. 2011; Chancerel *et al*. 2011). We therefore used a suite of approaches to maximise the representation of suitable SNPs on our array. Among the most important of these were (i) using multiple callers in our transcriptome variant discovery pipeline to identify a consensus SNP panel; (ii) mapping the flanking sequences of all SNPs to the fur seal reference genome to identify loci with the most suitable genomic contexts; and (iii) using Affymetrix design scores to filter out SNPs with unfavourable flanking sequence characteristics such as high GC content and non-specific hybridisation probabilities.

Overall, the comparably high conversion rate of our array suggests that these measures were successful. However, the total number of available SNPs was rather modest in relation to the size of the target array, meaning that we did not have a sufficient number of SNPs in our highest priority category to fill the entire array. Consequently, careful consideration was required when establishing additional prioritisation categories in order to strike a balance between SNP quantity and quality. In practice, we compromised on two main aspects. First, although we would have preferred only to tile loci with Affymetrix recommendations of “recommended”, this was not possible. Consequently, 37.9% of tiled SNPs had “neutral” Affymetrix recommendations. Second, Humble *et al*. (2018) found that loci mapping to more than one location in the reference genome were significantly less likely to convert, suggesting that probe sequence uniqueness may be an important factor to consider in SNP development. For this reason, we prioritised SNPs that mapped uniquely to the reference genome, although again we were constrained to include a number of SNPs whose flanking sequences revealed homology to more than one genomic region. As anticipated, conversion rates varied from a maximum of 93.7% for priority one SNPs down to a minimum of 66.3% for priority four SNPs. Interpreting these varying outcomes is not straightforward because SNPs in the various priority categories usually differed in multiple ways. Nonetheless, category one and two SNPs differed predominantly in their Affymetrix recommendations, so the 7% difference between these categories can be mainly attributed to this single factor.

Another strategy that we adopted to maximise genotyping success was to include SNPs that had been pre-validated using other technologies, including Illumina GoldenGate assays (Hoffman *et al*. 2012), KASP assays (Hoffman *et al*. 2013a) and Sanger sequencing (Humble *et al*. 2018). This approach was recommended by Kim et al. (2018), who reported higher rates of conversion on a 500K Affymetrix array for SNPs that had already been successfully genotyped on a 10K Illumina array. Unexpectedly, we found the opposite pattern, with pre-validated SNPs tending to perform worse on average than non-validated SNPs. The reasons for this remain unclear, although genotyping success was particularly low for SNPs derived from the canine HD bead chip. Our results therefore suggest that validating SNPs in advance may not always lead to better genotyping outcomes, especially when transferring loci from one technology to another.

As an alternative measure of genotyping success, we considered the proportion of samples that produced high quality genotypes. Only one fur seal sample out of 278 failed to pass quality control and three additional samples were considered to have failed because they fell a little short of the call rate threshold of 0.97. These numbers compare favourably with similar studies of both non-model organisms (e.g. Lundregan *et al*. 2018; Kim *et al*. 2018; Judkins *et al*. 2020). Overall, no relationship was found between the call rate per sample and DNA concentration, in contrast to Hagen *et al*. (2013) who reported that failed samples had significantly lower DNA concentrations than successful ones. However, all of our samples met or exceeded the recommended minimum total amount of DNA (200ng). Consequently, our findings are in agreement with (Kim *et al*. 2018), who experienced increased failure rates among samples that did not contain the recommended amount of DNA, but who found that DNA concentration did not influence genotyping success when sufficient amounts of DNA were provided.

### Levels of polymorphism

A very high proportion (97.3%) of the SNPs that successfully converted on the array were polymorphic in the Antarctic fur seal. Moreover, the true rate of polymorphism is probably higher, as several hundred SNPs were included on the array that showed *in silico* polymorphism in populations other than South Georgia, yet animals from these other localities were not genotyped on the array. Consequently, an unknown fraction of the SNPs that we have classified as monomorphic may in fact carry alleles that are private to one or more of the other populations. Our main reason for including these loci was to minimise ascertainment bias in future studies that might wish to genotype animals from different locations. Indeed, studies with similar discovery schemes have demonstrated negligible ascertainment bias towards populations from which the SNPs were initially discovered (van Bers *et al*. 2012; Malenfant *et al*. 2015; Kim *et al*. 2018).

Ascertainment bias cannot be avoided with SNP arrays because high frequency polymorphisms will always be easier to discover and can be called with greater confidence due to the minor allele being present in more individuals. Nevertheless, the strong positive association that we observed between *in silico* MAF and the empirical MAF of seals genotyped on the array suggests that, at least for moderately variable loci, the array provides a reasonable reflection of the underlying site frequency spectrum (SFS). This in turn suggests that the discovery pool of individuals in the original RAD sequencing study was large enough to estimate MAF reasonably well for the majority of SNPs that we built into the array. In line with this, a much weaker association was observed for the transcriptomic SNPs, which were discovered by sequencing many fewer individuals. Consequently, we do not recommend the array for approaches that may be sensitive to deviations from the true SFS, such as demographic inference. Nonetheless, for most purposes, SNPs with high MAFs are beneficial as they afford greater power for a multitude of applications ranging from parentage and relatedness analysis through linkage mapping to genome-wide association studies. Consequently, we believe this array will open up a wealth of new possibilities for delving into the population genomics of this important Antarctic predator.

### Inbreeding

To assess the levels of inbreeding in our study population we quantified three genomic inbreeding estimators (sMLH, 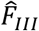 and *F*_ROH_). The resulting values were strongly intercorrelated, with *r* values ranging from 0.62 to 0.87, although associations involving *F*_ROH_ tended to be weaker. When using incomplete marker information from a SNP chip, short ROH arising from inbreeding in the very distant past cannot be reliably detected due to inadequate SNP densities (Kardos *et al*. 2016). To take account for this, we only called ROH segments that were above a stringent length threshold. Furthermore, to avoid spurious ROH calls caused by low marker densities, we only considered ROH segments present in regions of the genome represented by high marker densities. Therefore, whilst our measures of sMLH and 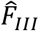 have captured variation in inbreeding due to IBD segments arising from both recent and distant ancestors, our measure of *F*_ROH_ is unlikely to have captured variation in inbreeding due to very distant ancestors. Additionally, our estimates of *F*_ROH_ might be less reliable in the current study due to the fragmented nature of our reference genome, which could potentially have introduced noise into our estimates. However, if anything, we expect these factors to have led to the magnitude of *F*_ROH_ being underestimated. We hope to be able further refine our estimates of inbreeding in future studies by improving the contiguity of the fur seal reference genome and by calibrating array-based measures of inbreeding by reference to whole genome resequencing data.

Nevertheless, the fact that *F*_ROH_ was non-zero in all of our samples despite the conservative nature of our analysis provides support for the presence of inbreeding in the study population. Most individuals carried ROH segments making up around 6% of the genome, with *F*_ROH_ ranging from as little as 2% in one individual to as much as 8% in four individuals. These numbers are comparable with estimates for other wild mammal populations such as the Iberian ibex (Grossen *et al*. 2018), Dryas monkey (van der Valk *et al*. 2020) and Icelandic horse (Schurink *et al*. 2019), and suggest that previously documented correlations between heterozygosity and fitness may be due to inbreeding depression (Hoffman *et al*. 2004, 2007; Forcada and Hoffman 2014). Furthermore, the vast majority of ROH segments were shorter than 5 Mb, with only four individuals harbouring ROH longer than 20 Mb. Therefore, most of the IBD observed in our study population has probably arisen from inbreeding between ancestors in the distant past, as opposed to inbreeding in more recent generations. These findings suggest that the population of Antarctic fur seals is large enough to minimise very close inbreeding and / or that female mate choice is effective in preventing matings between close relatives (Hoffman *et al*. 2007).

### Relatedness

Our study also illustrates the potential for high density SNP genotype data to recover known relationships and to uncover the relatedness structure of a sample of individuals. Genome-wide measures of relatedness based on IBD allele sharing confidently identified the positive controls and were also able to flag the presence of known mother-offspring pairs in our dataset. Nevertheless, we found that field-based assignments of mothers to pups were not always correct, in support of a previous study that found high rates of fostering and milk-stealing in the study colony (Hoffman and Amos 2005). We were initially surprised to discover over 60 pairs of related individuals (0.1 ≤ *r* ≤ 0.3) in our sample. Investigating this in greater detail, we uncovered evidence in support of the presence of a mixture of second order relatives (which could potentially include additional half siblings, avuncular and grandparent-grandchild relationships), and third order relationships (such as possible first cousins). Notably, full siblings were conspicuously absent from our dataset, in contrast to grey seals, where around 30% of offspring are full siblings due to partner fidelity (Amos *et al*. 1995). However, mate fidelity is unlikely to be very important in Antarctic fur seals because the vast majority of territorial males only come ashore for one or two seasons in total (Hoffman *et al*. 2003).

Unfortunately, we were not able to ascertain the exact nature of the majority of cryptic relationships within our dataset because sequoia was constrained by a lack of known pedigree links other than mother-offspring pairs. Nevertheless, sequoia confidently identified four pairs of paternal half siblings, which we would expect to be present in the study colony given the polygynous mating system of this species (Hoffman *et al*. 2003). To shed further light on the relatedness structure of the study colony would require the construction of a multigenerational pedigree. In the past, we have considered this problematic due to the long generation time of this species relative to the duration of our study and the fact that not all the pups are sampled every year. However, the potential for augmenting classical microsatellite based parentage analysis with genomic information gives us new grounds for optimism.

### Cross-species amplification

Finally, we explored the cross-species amplification potential of our array by genotyping small numbers of grey seals, Galápagos sea lions and Steller’s sea lions. Although none of the grey seals passed quality control, over 70,000 loci cross-amplified in both of the otariid species and over five percent of these were polymorphic, yielding over 4,000 polymorphic SNPs per species. This is in line with expectations set out in Miller *et al*. (2015) and demonstrates the applicability of the array for generating genomic data in closely related pinniped species. It may also be worth considering testing the array on less divergent pinnipeds, most obviously other fur seal species belonging to the genus *Arctocephalus*, some of which diverged from *A. gazella* as recently as around one million years ago (Higdon *et al*. 2007) and where rates of polymorphism are expected to be as high as 20–90% (Miller *et al*. 2012).

### Conclusions

SNP arrays provide a straightforward and effective solution for generating very large genetic marker datasets encompassing many individuals. As such, they have been instrumental in opening up a wide variety of questions to investigation in natural populations, from population genomics to quantitative genetics. This manuscript describes the successful development and implementation of a SNP array for a model marine mammal species, the Antarctic fur seal. By employing strict filtering approaches incorporating knowledge of the genomic context of each SNP, we were able to achieve comparably high rates of conversion and polymorphism. We also confirmed and built upon the results of previous studies by quantifying both inbreeding and genomic relatedness. We hope not only that our array will open up new avenues in fur seal research, but also that the protocols we developed to improve genotyping outcomes will be applicable to the design of arrays for other species.

## ACKNOWLEDGEMENTS

This research was supported by the Deutsche Forschungsgemeinschaft (DFG, German Research Foundation) in the framework of a Sonderforschungsbereich (project numbers 316099922 and 396774617–TRR 212) and the priority programme “Antarctic Research with Comparative Investigations in Arctic Ice Areas” SPP 1158 (project number 424119118). It was also supported by core funding from the Natural Environment Research Council to the British Antarctic Survey’s Ecosystems Program. We are grateful to the many field assistants on Bird Island who contributed toward animal handling and tissue sampling. Samples were collected as part of the Polar Science for Planet Earth program of the British Antarctic Survey under the authorisation of the Senior Executive and the Environment Officers of the Government of South Georgia and the South Sandwich Islands. Samples were collected and retained under Scientific Research Permits for the British Antarctic Survey field activities on South Georgia, and in accordance with the Convention on International Trade in Endangered Species of Wild Fauna and Flora. All field procedures were approved by the British Antarctic Survey Animal Welfare and Ethics Review Body.

## AUTHOR CONTRIBUTIONS

JIH and EH conceived and designed the study. JF and JIH contributed materials and funding. EH and AJP prepared the samples for genotyping. EH designed the SNP array and analysed the data. EH and JIH wrote the manuscript. All of the authors commented on and approved the final manuscript.

